# Human POMC processing *in vitro* and *in vivo* revealed by quantitative peptidomics

**DOI:** 10.1101/257121

**Authors:** Peter Kirwan, Richard Kay, Bas Brouwers, Vicente Herranz-Perez, Magdalena Jura, Pierre Larraufie, Jason Pembroke, Theresa Bartels, Anne White, Fiona Gribble, Frank Reimann, I. Sadaf Farooqi, Stephen O’Rahilly, Florian T. Merkle

**Author notes:** To whom correspondence should be addressed, WT-MRC Institute of Metabolic Science (Level 4), Addenbrooke’s Site (B289), Keith Day Road, Cambridge, CB2 0QQ, UK.

## Abstract

Human obesity can result from the aberrant production or processing of proopiomelanocortin (POMC) in hypothalamic neurons, but it is unclear which human POMC-derived peptides are most relevant to body weight regulation. To address this question, we analysed both hypothalamic neurons derived from human pluripotent stem cells (hPSCs) and primary human hypothalamic tissue using quantitative liquid chromatography tandem mass spectroscopy (LC-MS/MS). In both *in vitro-* and *in vivo*-derived samples, we found that POMC was processed into β-melanocyte stimulating hormone (β-MSH), whose existence in the human brain has been controversial. β-MSH and desacetyl α-MSH (d-α-MSH) were produced at roughly equimolar concentrations and in vast excess to acetylated α-MSH (5-to 200-fold), suggesting that the importance of both d-α-MSH and β-MSH to human obesity has been underestimated. Since body weight is sensitive to changes in MSH concentration, we asked whether hPSC-derived hypothalamic neurons could provide mechanistic insights into the processing and secretion of MSH peptides. We found that cultured human hypothalamic neurons appropriately trafficked POMC and its derivatives, and robustly (P<0.0001) secreted them when depolarised. Furthermore, the adipocyte-derived hormone leptin significantly (P<0.01) promoted their production of both d-α-MSH and β-MSH. These results establish hPSC-derived hypothalamic neurons as a model system for studying human-specific aspects of POMC processing that might be therapeutically harnessed to treat obesity.

## INTRODUCTION

Obesity is a major public health problem. In order to develop effective treatments, it is important to understand the endogenous mechanisms that regulate energy homeostasis. Hypothalamic neurons that produce proopiomelanocortin (POMC) are important regulators of energy homeostasis in both mice^1^ and humans^2^. POMC protein undergoes extensive proteolytic cleavage to produce neuropeptides that regulate food intake and energy expenditure^3,4^ (Fig. 1). These peptides include desacetyl α-melanocyte stimulating hormone (ACTH(1-13) amide, d-α-MSH(1-13), d-α-MSH), which can also be acetylated to produce a-MSH(1-13). Both d-α-MSH and α-MSH stimulate melanocortin receptor 4 (MC4R) in the brain^5^^-^^8^ to reduce food intake. POMC is also processed into β-endorphin (β-EP(1-31))^9,10^ which stimulates the µ-opioid receptor to promote analgesia and increased food intake^11^^-^^13^. Human hypothalamic POMC neurons might also produce β-MSH^14^, which is associated with human obesity when mutated^15^^-^^17^ and inhibits feeding as potently as α-MSH when injected into mice^17^ by activating MC4R^16^^-^^18^. However, β-MSH is not produced in the rodent brain and its existence and biological relevance in humans is controversial^19^^-^^21^.

**Figure 1:**
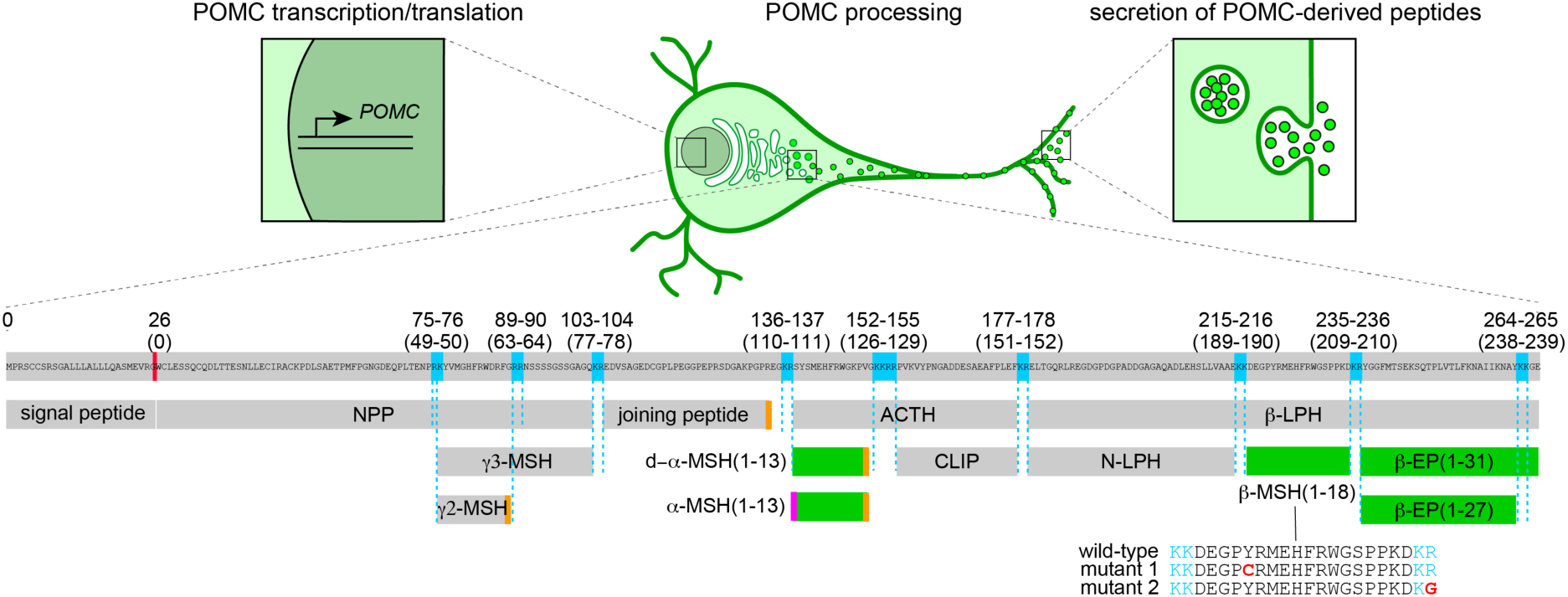
Regulation of POMC processing and secretion. Human POMC is translated as a 267-amino acid protein that, after removal of the signal peptide, undergoes successive rounds of cleavage and trimming at dibasic residues (blue) in a tissue-specific manner; the hypothalamic pattern is illustrated. Additional levels of post-translational modification include C-terminal amidation (orange) and N-terminal acetylation (magenta). POMC-derived peptides that regulate food intake (green) include d-α-MSH(1-13), α-MSH(1-13), β-MSH(1-18), and β-endorphin (β-EP(1-31) and β-EP(1-27)). Illustrated mutations in β-MSH have been associated with obesity, suggesting a role for this peptide in human body weight regulation. The concentrations of secreted POMC-derived peptides may be regulated at the levels of transcription, translation, processing, and secretion. ACTH, adrenocorticotropic hormone; CLIP, corticotropin-like intermediary peptide; EP, endorphin; LPH, lipotropin; MSH, melanocyte stimulating hormone; NPP, POMC N-terminal region; POMC, proopiomelanocortin.

Since body weight is sensitive to changes in the concentration of POMC-derived peptides^22-26^, understanding POMC processing can provide insights into molecular mechanisms of obesity that might be therapeutically harnessed^3^. Indeed, obesity results from the reduced functionality of the POMC-processing enzymes proprotein convertase subtilisin/kexin type 1 (*PCSK1, PC1/3*)^27,28^, carboxypeptidase E (*CPE*)^29,30^, or the transcription factor nescient helix-loop-helix 2 (*NHLH2*) that regulates *PCSK1* expression^31,32^. Since POMC processing is incomplete even in wild-type brains^9,10^, stimulating POMC production or promoting its processing might reduce body weight in obesity. For example, the adipocyte-derived hormone leptin promotes POMC gene expression^25,33^ and POMC processing^34,35^ to increase α-MSH production in mice.

While studies in mice have been valuable, rodents do not produce β-MSH, limiting their utility as a model system for studying human-specific aspects of POMC processing. Postmortem human brain samples provide a snapshot of peptide production, but cannot provide information on the dynamic regulation of POMC processing and secretion. To address these issues, we^36,37^ and others^38,39^ developed methods to generate hypothalamic neurons in culture from human pluripotent stem cells (hPSCs). Since these hypothalamic neurons can be produced at scale and are readily manipulated, they provide an unprecedented opportunity to study human POMC biology. POMC-derived peptides are typically quantified using ELISAs, but these assays only provide information about particular target peptide and often cannot discriminate between peptides that share common epitopes but have functionally relevant structural differences^10,40-42^. In contrast, liquid chromatography tandem mass spectroscopy (LC-MS/MS) is sensitive and provides information on the precise sequence and post-translational modifications of thousands of peptides in a single experiment^43^.

Here, we used LC-MS/MS to analyse POMC processing in both hPSC-derived hypothalamic neurons and primary human brain samples to address several outstanding questions: *Is β-MSH produced in the human brain? What is the relative abundance of POMC-derived peptides that regulate human energy homeostasis? Is POMC processing in human hypothalamic neurons regulated by exogenous factors such as leptin?*

We found that hPSC-derived hypothalamic neurons trafficked and secreted POMC and its derivatives, and appropriately processed POMC into neuropeptides previously established to regulate energy balance, both *in vitro* and *in vivo.* These neuropeptides included β-MSH(1-18). We next developed quantitative assays for POMC-derived peptides and found that acetylated α-MSH was present at substantially (5- to 200- fold) lower concentrations than d-α-MSH, β-MSH, or β-EP. The production of d-α-MSH and β-MSH could be significantly (P<0.01) stimulated by treatment with the adipocyte-derived hormone leptin, demonstrating that POMC processing can be dynamically regulated *in vitro*. Together, these studies establish hPSC-derived hypothalamic neurons as a valuable tool for studying the dynamics of POMC processing and secretion, and suggest that the roles of β-MSH and d-α-MSH in human obesity have been underappreciated.

## RESULTS

### Human hypothalamic neurons appropriately traffic and secrete POMC *in vitro*

To determine if *in vitro*-derived human POMC neurons could be a suitable model system for studying POMC processing, we differentiated human embryonic stem cells (hESCs) and human induced pluripotent stem cells (hiPSCs) into hypothalamic neurons as previously described^36,37^ (Fig. 2A). After 28 days of differentiation, cultures expressed hypothalamic marker genes such as NKX2.1, and predominantly contained cells with neuronal morphology that expressed the neuronally-enriched proteins tubulin beta 3 class III (TUBB3, Tuj1) and microtubule associated protein 2 (MAP2) (Fig. 2B). We observed that 6.5 ± 0.76 percent of neurons were strongly immunoreactive for POMC, as visualised by several distinct antibodies (Fig. 2B-Fig. 2D).

**Figure 2:**
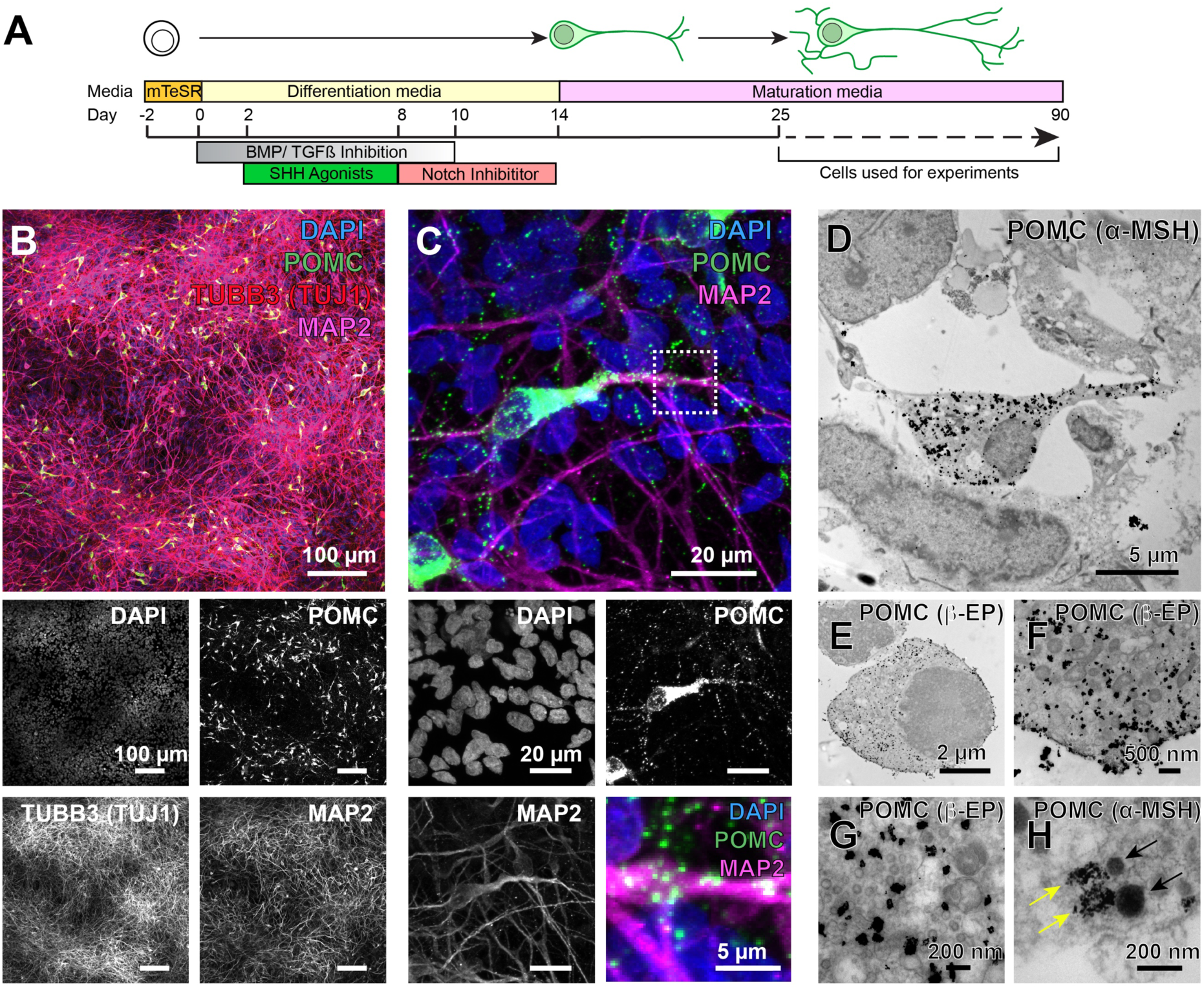
Subcellular localisation of POMC and its derivatives in hPSC-derived hypothalamic neurons. **A**) Schematic diagram of hypothalamic differentiation from human pluripotent stem cells (white) into hypothalamic POMC neurons (green) that mature over time in culture. Experiments were carried out between 25-90 days post-differentiation. **B**) Human hypothalamic differentiation yielded predominantly neuronal cultures as indicated by immunostaining for TUBB3 (Tuj1) and MAP2, of which approximately 6% were immunopositive for POMC. **C**) High-magnification confocal microscopy of hPSC-derived hypothalamic neurons revealed that POMC-immunoreactive structures were punctate and localised to cell bodies and neurites, consistent with a vesicular distribution pattern (see higher magnification inset). **D,E**) Transmission electron microscopy of fixed hPSC-derived hypothalamic neurons stained with Immunogold for α-MSH (D) or β-EP (E) revealed that labelling was widespread in the cytoplasm and neurites, excluded from the nucleus, and absent from surrounding neurons. **F,G**) High magnification TEM showing Immunogold labelling localised to rounded, vesicle-like structures in regions with abundant vesicles. **H**) α-MSH-labelled structures (yellow arrows) were often found adjacent to dense core vesicles (black arrows).

Since POMC becomes packaged into dense core vesicles as it is progressively processed^44^^-^^47^, we examined subcellular localization of POMC-immunoreactive epitopes by high-magnification immunofluorescent confocal microscopy and found that they localised to punctate structures within the cell body and neurites (Fig. 2C). To ask if these structures corresponded to dense-core vesicles, we performed Immunogold labelling with antibodies recognising epitopes corresponding to sequences in α-MSH or β-EP and examined samples by transmission electron microscopy (TEM). Immunogold particles localised (Fig. 2D-Fig. 2H) to rounded structures approximately 50-250 nm in diameter (Fig. 2F,Fig. 2G), some of which resembled and were adjacent to dense-core vesicles (Fig. 2H). These results suggest that POMC or its derivative peptides are packaged into dense-core vesicles.

Since these findings suggested that POMC might be secreted, we analysed the culture media collected from hPSC-derived cortical or hypothalamic cells using a sensitive (detection limit ~5 pM) sandwich ELISA. This assay detects epitopes near ACTH(10-18) and NPP^41,42,48^, both of which are present in full-length POMC and in pro-ACTH (Fig. 1, S1 A,B). We could readily detect these species in media from hypothalamic cultures but not from cortical cultures (Fig. S1C), consistent with their constitutive secretion and/or secretion in response to spontaneous neuronal activity present during standard culture conditions. To test if secretion might be stimulated by depolarisation, we treated cultures with KCl (Fig. S1D) and observed a significant (P<0.01) 6.3 ± 0.16-fold increase in secreted POMC and/or pro-ACTH (Fig. S1E).

**Figure S1:**
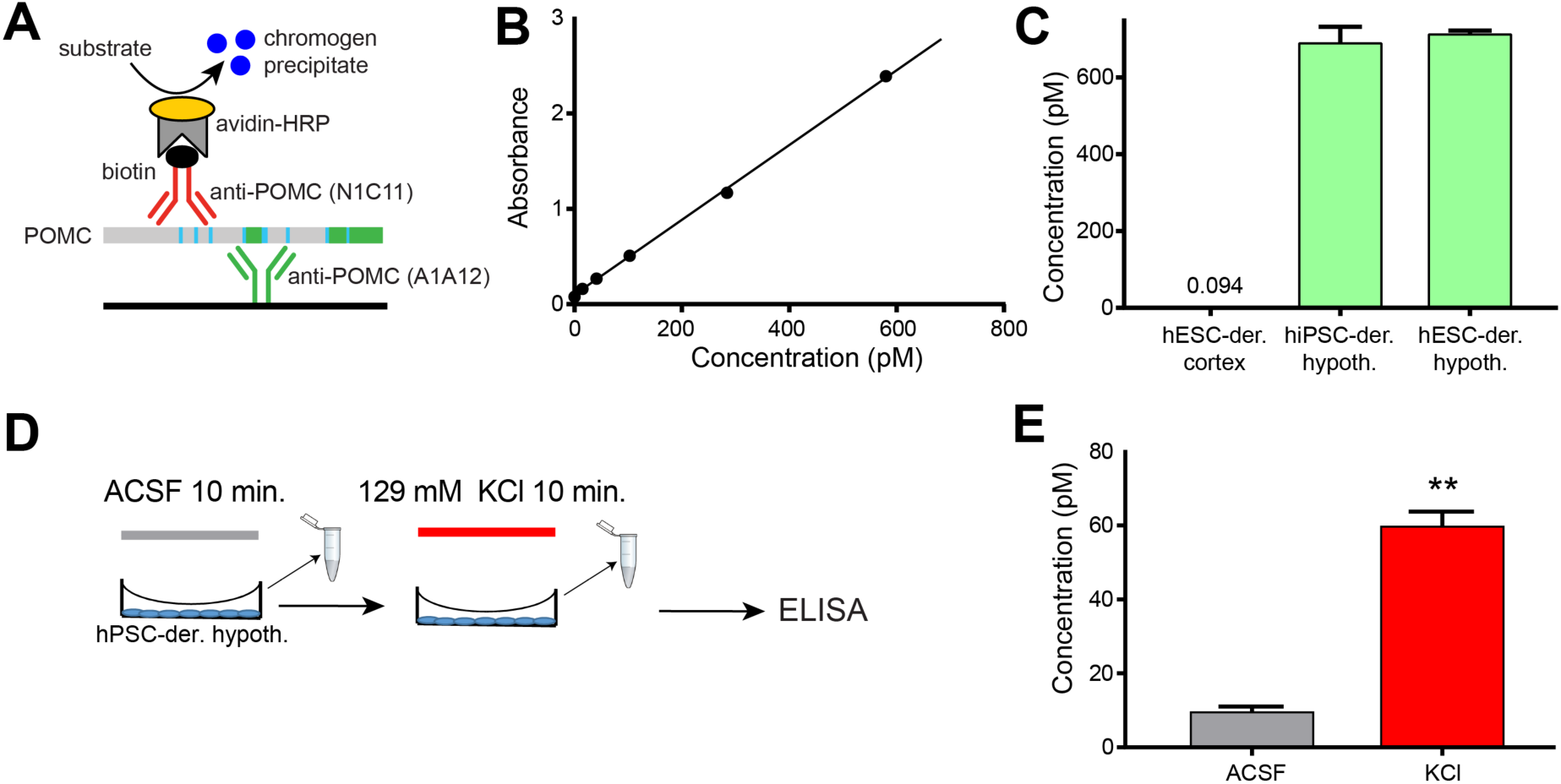
Secretion of POMC in hPSC-derived hypothalamic neurons in the presence or absence of experimental depolarisation. **A**) Schematic diagram of the sandwich ELISA assay for detecting full-length POMC using monoclonal anti-POMC antibodies that detect epitopes near ACTH(10-18) and NPP, both of which are present in full-length POMC and pro-ACTH, enabling absorbance-based quantification. **B**) The ELISA standard curve generated using recombinant POMC protein is linear over a broad range and enables quantification down to approximately 5 pM. **C**) Full-length POMC and/or pro-ACTH were not detected within the quantification limits of the assay from hPSCs differentiated to cortical neurons, but were robustly detected from either hiPSCs or hESCs differentiated to hypothalamic neurons. N=2 independent cell lines, 3 replicate wells per cell line **D**) Schematic diagram of stimulation experiments to test for depolarisation-induced increases in POMC and/or pro-ACTH secretion by ELISA. **E**) KCl-induced depolarisation significantly (P<0.01) increased measured concentrations of secreted POMC and/or pro-ACTH. N=1 experiment, 3 replicate wells. ACSF, artificial cerebrospinal fluid; der., derived; HRP, horseradish peroxidase; hypoth., hypothalamic, KCl, potassium chloride; **, P <0.01. Error bars show SEM.

### hPSC-derived hypothalamic neurons produce b-MSH(1-18) and other POMC-derived peptides

To identify which POMC-derived peptides were produced in human hypothalamic neurons, we extracted peptides using either 80% acetonitrile (ACN) or 6M guanidine hydrochloride (GuHCl) followed by ACN, and analysed them by nanoflow LC-MS/MS^49^ (Fig. 3A). We observed similar types of peptides using these two extraction methods, but found that GuHCl treatment increased both the number of unique peptides and the depth of spectra observed per peptide (Table S1) (Personal communication,
Pierre Larraufie et al.).

**Figure 3:**
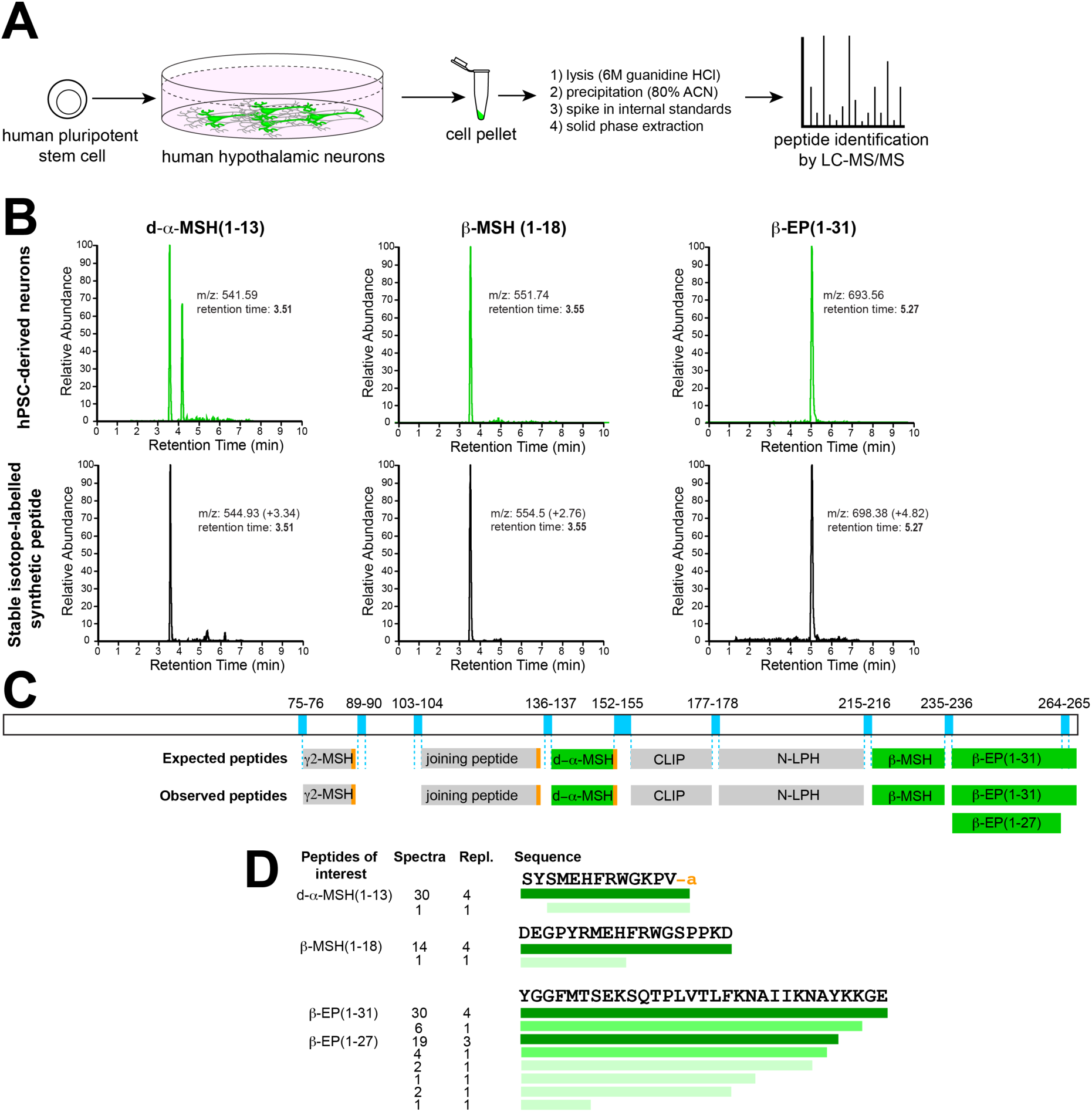
Identification of processed POMC peptides by LC MS/MS: **A**) Schematic workflow for the quantification of POMC-derived peptides from hPSC-derived hypothalamic neurons. **B**) Liquid chromatographs of samples from hPSC-derived neurons (top), compared with stable isotope-labelled synthetic d-α-MSH(1-13), β-MSH(1-18) and β-EP(1-31) (bottom). Note the identical retention times for endogenous and reference peptides, which had slightly higher mass-to-charge (m/z) ratios due to the incorporation of heavy isotopes. Note that Q1 m/z values measured from the Orbitrap and triple quadripole mass spectrophotometers may be differ. **C**) Schematic of human POMC protein and relevant dibasic cleavage sites (blue), expected POMC-derived peptides, and those peptides detected in hPSC-derived hypothalamic cultures by LC-MS/MS. Quantified peptides are indicated in green. **D**) Summary of POMC-derived peptides detected by LC-MS/MS, where the colour intensity represents the relative abundance of each peptide species. Note that although some degradation products are present, the predominant forms observed *in vitro* match those expected from previous studies performed *in vivo*. Repl., replicates. Other abbreviations are as in Fig. 1.

**Table S1:** Detection of neuropeptides by LC-MS/MS. Peptide extraction by either 80% ACN alone or 6M GuHCl followed by ACN yielded similar types of peptides, but GuHCl-ACN extraction was associated with a larger number of peptides and a larger number of spectra supporting each peptide call.

| **Protein** | **80% ACN** |  | **GuHCl-ACN** |  |
| --- | --- | --- | --- | --- |
|  | **# Peptides** | **# Spectra** | **# Peptides** | **# Spectra** |
| Calcitonin | 0 | 0 | 1 | 1 |
| Chromogranin A | 6 | 12 | 10 | 33 |
| Neurosecretory Protein VGF | 9 | 20 | 75 | 372 |
| Neurotensin | 0 | 0 | 1 | 6 |
| Secretogranin 1 | 7 | 19 | 12 | 38 |
| Secretogranin 2 | 4 | 11 | 9 | 31 |
| Secretogranin 3 | 0 | 0 | 4 | 14 |
| Thymosinβ10 | 24 | 123 | 40 | 284 |
| Thymosinβ4 | 33 | 120 | 45 | 249 |

To test whether hPSC-derived hypothalamic cultures processed POMC into peptides important for energy homeostasis, we custom-synthesised d-α-MSH(1-13), β-MSH(1-18) and β-EP(1-31) with specific amino acids substituted for ^13^C and ^15^N stable isotopically labelled equivalents, and spiked these internal standards into cell lysates during peptide extraction (Fig. 3A) (see also Materials and Methods). Stable isotope-labelled standards have shifted mass-to-charge ratios (m/z) compared with endogenous peptides, but are otherwise chemically identical, resulting in identical column retention times during liquid chromatography (Fig. 3B).

We then identified peptides present in samples from two unrelated hESC lines and one hiPSC line using an automated peptide identification pipeline (see Materials and Methods). We consistently observed POMC-derived peptides in hPSC-derived hypothalamic cultures (but not in cortical cultures), constituting 3.35 ± 1.87 % of all peptide alignments (Table S2). The vast majority of unique spectra for POMC-derived peptides of interest (50/69, 72.4%) were flanked by dibasic residues that are targeted by POMC-processing enzymes (Fig. 3C), and some peptides were C-terminally amidated as is seen *in vivo* (31/31 analysed spectra), suggesting that the observed peptides were unlikely to be random degradation products (Fig. 3D). We identified the POMC-derived neuropeptides d-α-MSH, β-MSH and β-EP(1-31) as well as β-EP(1-27), g2-MSH, CLIP, N-LPH, and joining peptide (Fig. 1, Fig. 3C). These results extend a previous report^50^ by providing the specific sequences and post-translational modification patterns of these peptides and establishing that β-MSH(1-18) is produced by human hypothalamic neurons.

**Table S2:** Quantitative summary of the number of spectra, peptides and proteins in hPSC-derived hypothalamic neurons (*in vitro*) as well as from primary human brain (*in vivo*). A summary of the number of POMC-derived peptides and supporting spectra is also shown, as well as the percentage for POMC spectra and peptides as a function of total peptides observed in each experiment.

| ***In vitro*** |  |  |  |  |  |  |
| --- | --- | --- | --- | --- | --- | --- |
| Sample | Sample 1 (H9 ES Day 84) | Sample 2 (H9 ES Day 84) | Sample 3 (13B iPS Day 90) | Sample 4 (HUES9 ES Day 40) | Mean | SEM |
| Extraction method | ACN | ACN | ACN | GuHCl-ACN |  |  |
| Spectra | 7448 | 7217 | 423 | 4218 | 4826.50 | 1641.84 |
| Peptides | 3436 | 3031 | 358 | 1830 | 2163.75 | 691.79 |
| Proteins | 967 | 757 | 204 | 427 | 588.75 | 169.70 |
| POMC spectra | 26 | 168 | 8 | 372 | 143.50 | 84.15 |
| % POMC spec/total spec | 0.35 | 2.33 | 1.89 | 8.82 | 3.35 | 1.87 |
| POMC peptides | 9 | 40 | 7 | 55 | 27.75 | 11.81 |
| % POMC pep/total pep | 0.26 | 1.32 | 1.96 | 3.01 | 1.64 | 0.57 |
| ***In vivo*** |  |  |  |  |  |  |
| Sample | Cortex | Mediobasal hypothalamus | Paraventricular hypothalamus | Mean | SEM |  |
| Extraction method | GuHCl-ACN | GuHCl-ACN | GuHCl-ACN |  |  |  |
| Spectra | 1018 | 8971 | 3427 | 4472 | 2354.54 |  |
| Peptides | 732 | 3666 | 1871 | 2089.67 | 854 |  |
| Proteins | 210 | 481 | 454 | 381.67 | 86.19 |  |
| POMC spectra | 0 | 24 | 3 | 9 |  |  |
| % POMC spec/total spec | 0.00 | 0.27 | 0.09 |  |  |  |
| POMC peptides | 0 | 7 | 3 |  |  |  |
| % POMC pep/total pep | 0.00 | 0.19 | 0.16 |  |  |  |

We were surprised to readily detect spectra for β-MSH(1-18) whose existence has been disputed^20^, whereas we did not consistently detect spectra for α-MSH(1-13) which has been the subject of intense study^6,7,35,51,52^ since it is produced in mice and suggested by some studies to be more stable than d-α-MSH^35^. Upon manual review, we could indeed detect peaks with m/z and column retention times appropriate for α-MSH. We therefore wondered what the relative concentrations of POMC-derived peptides relevant for human energy homeostasis might
be *in vitro* and *in vivo*.

### d-α-MSH(1-13), β-MSH(1-18), and β-EP(1-31) are produced in excess of α-MSH(1-13)

To address this question, we developed quantitative assays^53^ for α-MSH(1-13), d-α-MSH(1-13), β-MSH(1-18) and β-EP(1-31) (Fig. 4A). We first generated calibration lines (R^2^>0.98) with synthetically produced human peptides (Fig. 4B, 4C), enabling accurate quantification down to at least 10 pg/ml (2.9 to 6 pM). Calibration lines were then compared to those obtained from samples spiked with internal standard peptides, enabling endogenous peptides to be accurately quantified across a broad concentration range. Since α-MSH and d-α-MSH are structurally and chemically similar, and to facilitate direct comparison of endogenous α-MSH and d-α-MSH to a single standard within the same sample, we compared both of these peptides to an internal standard for d-α-MSH.

**Figure 4:**
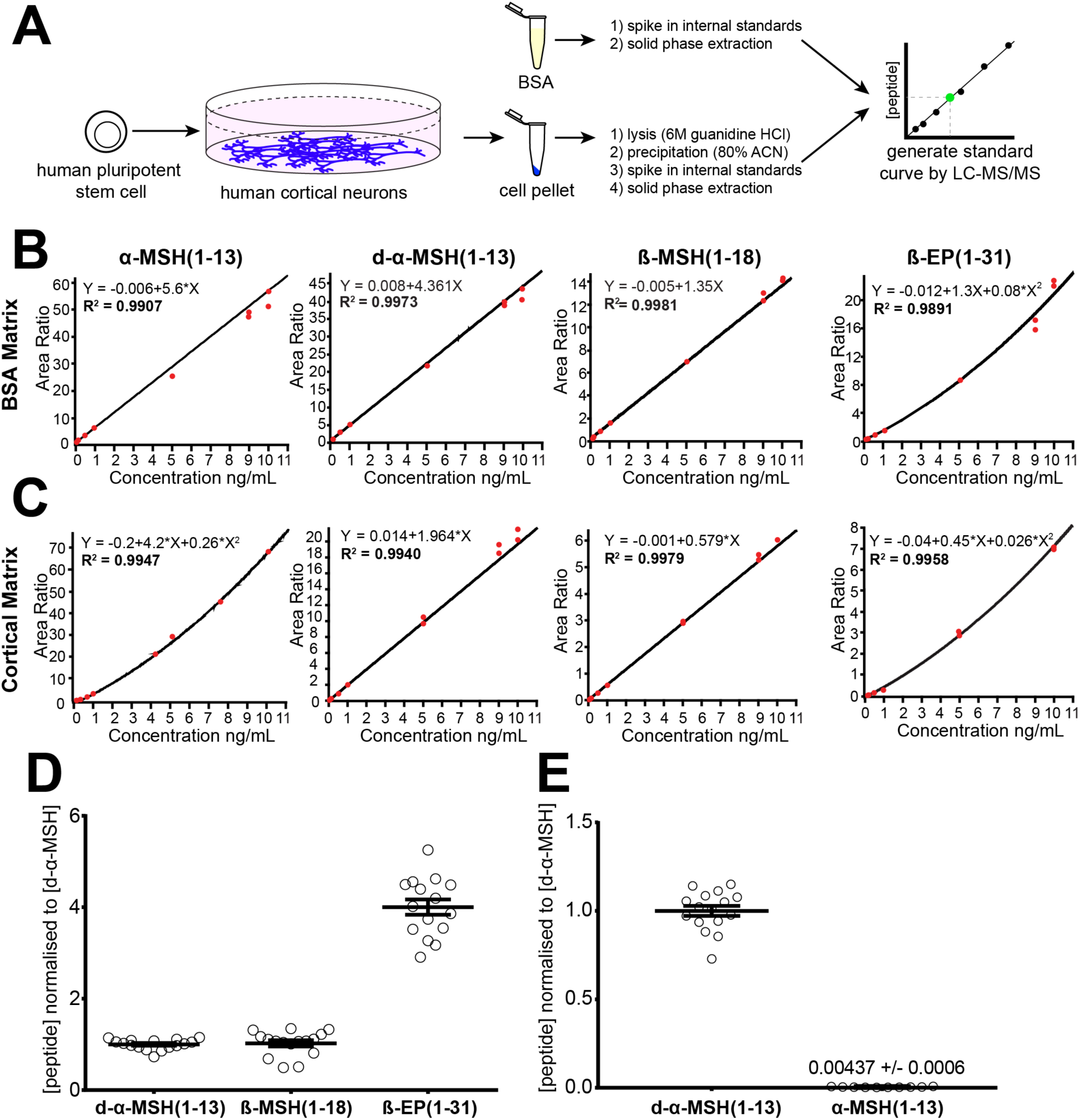
Quantification of POMC-derived peptides in hPSC-derived neurons. **A**) Schematic diagram of peptide standard curve generation. **B,C**) Standard curves were generated by adding known quantities of synthetic POMC-derived peptides to a BSA matrix (B) or hPSC-derived cortical cell matrix (C) enable peptide quantification (R^2^>0.98) over a broad concentration range. **D,E**) LC-MS/MS-based quantification of d-αMSH, β-MSH and β-EP peptides in hPSC-derived POMC neurons (D) and direct comparison of d-α-MSH and α-MSH concentrations within samples. Concentrations were converted to molarity and then normalised to d-α-MSH. N=4 independent experiments with 3-6 replicates per experiment. Error bars show SEM.

We found that target peptides were typically present at quantities greater than 540 pM, well above the quantification limit of the assay. Upon comparing the relative concentrations of these peptides within a particular sample of hPSC-derived hypothalamic neurons across four independent experiments, we found that d-α-MSH and β-MSH were present at equimolar ratios (1 ± 0.028 d-α-MSH: 1.027 ± 0.067 β-MSH), and that β-EP(1-31) concentrations were 4 ± 0.167- fold higher than either of these two MSH species (Fig. 4D). In contrast, α-MSH was often present at concentrations near the assay’s detection limit.Whether we considered all data for α-MSH or limited our analysis to samples with concentrations above 6 pM, we observed that α-MSH was present at less than one hundredth the concentration of d-α-MSH at the developmental time points analysed (Fig. 4E).

The analysis of hPSC-derived hypothalamic cultures yielded many results consistent with what has been described in other experimental systems, but we were surprised by the relative abundance of both d-α-MSH and β-MSH relative to α-MSH. To test whether observations made *in vitro* were predictive of the concentrations of these peptides *in vivo*, we obtained two fresh post-mortem human brain samples and dissected regions encompassing the mediobasal hypothalamus (MBH), the paraventricular nucleus of the hypothalamus (PVH), and a small piece of the orbitofrontal gyrus of the cerebral cortex (CTX) from each brain (Fig. 5A,B).

**Figure 5:**
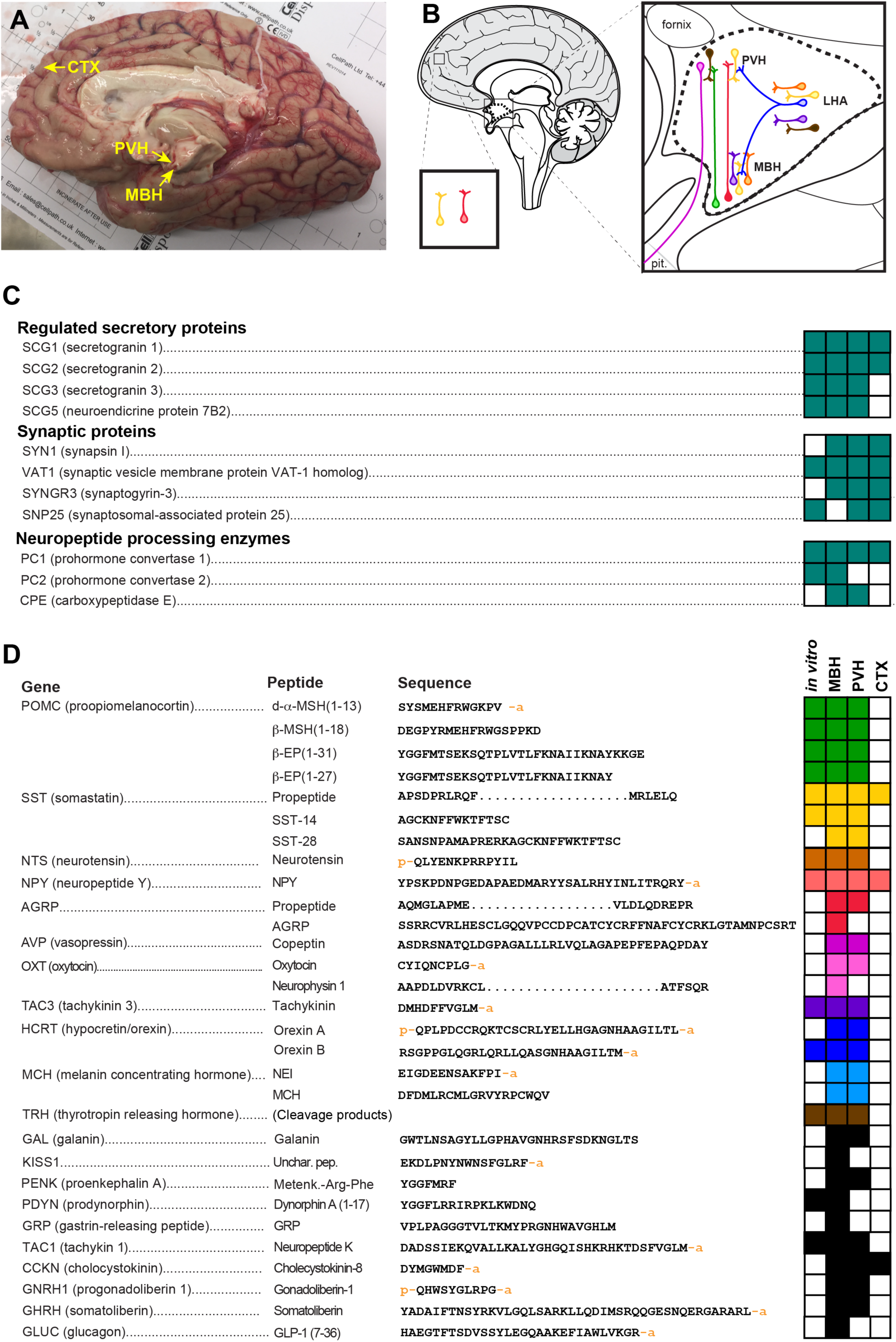
Characterisation of the peptidome of the human hypothalamus. **A**) Sagittal view of the right hemisphere of a post-mortem human forebrain used in this study. Regions of approximately 3x3x3 mm from the indicated brain regions were dissected for analysis. **B**) Colour-coded diagram of several neuropeptidergic cell types of interest, and some of their projections. **C**) Summary of peptides associated with the regulated secretory pathway, synaptic proteins, and neuropeptide processing enzymes detected in hPSC-derived hypothalamic neurons *in vitro* or in the MBH, PVH, or CTX of the human brain. Filled boxes represent that peptides corresponding to the indicated gene were detected and automatically identified. **D**) Summary of select neuropeptides detected in hPSC-derived hypothalamic neurons or in the human brain. Sequences too long to readily display here are denoted by an ellipsis. CTX, cortex (orbitofrontal gyrus); LHA, lateral hypothalamic area; MBH, mediobasal hypothalamus; Pit., pituitary gland; PVH, paraventricular nucleus of the hypothalamus. An adjoining ‘p’ on the amino acid sequences denotes a pyroglutamate residue, while an ‘a’ denotes an amide group.

By nanoflow LC-MS/MS analysis of these brain samples, we identified an average of 2089 ± 854 peptides from 4472 ± 2355 spectra per sample, corresponding to 382 ± 86 unique gene products (Table S2). These results are comparable to what was observed in hPSC-derived hypothalamic neuronal cultures (2164 ± 691 peptides, 4827 ± 1642 spectra, 588 ± 170 gene products per sample) (Table S2). In each primary sample we observed peptides associated with synapses and regulated secretion, suggesting that peptides were effectively extracted from each brain region. In primary human hypothalamic samples, we detected several key enzymes involved in neuropeptide processing (Fig. 5C). Primary human MBH and PVH samples contained the POMC-derived peptides α-MSH(1-13), d-α-MSH(1-13), β-MSH(1-18) and β-EP(1-31), along with a broad array of other neuropeptides such as agouti-related peptide (AGRP) (Fig. 5D). Many of the peptides identified in the human brain were also present with identical structures and post-translational modifications in
hPSC-derived hypothalamic neurons (Fig. 5D).

To determine the relative concentrations of POMC-derived peptides in the human brain, we used the quantification methods based on stable-isotope-labelled internal standards as described above. In agreement our observations in hPSC-derived hypothalamic cultures, we found that β-MSH was indeed present in the adult human MBH and PVH at concentrations comparable to that of d-α-MSH. We also found that β-EP(1-31) was present at approximately 2- to 3-fold higher concentrations than either d-α-MSH or β-MSH but that acetylated α-MSH concentrations were approximately 6- to 10-fold lower than those of either d-α-MSH or β-MSH (Fig. S2). These results were largely consistent with those seen in hPSC-derived hypothalamic neurons, suggesting that these *invitro* cellular models might provide insights into the dynamic regulation of POMC expression, processing, and secretion.

**Figure S2:**
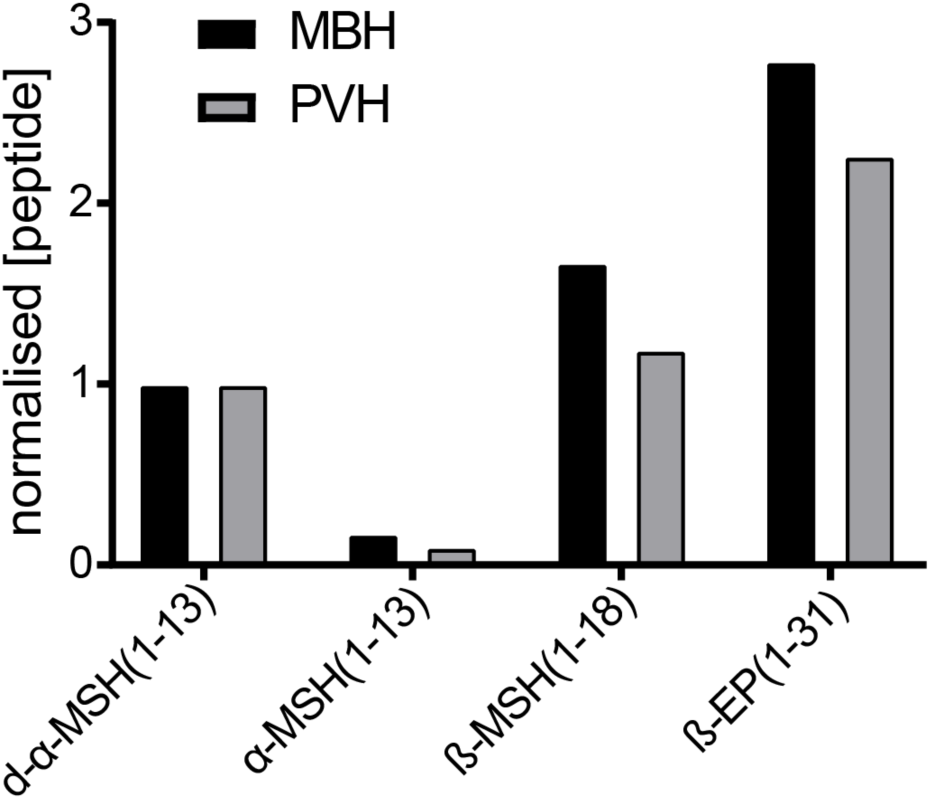
d-α-MSH(1-13) and β-MSH(1-18) are more abundant than α-MSH(1-13) in the human hypothalamus. Quantitative LC-MS/MS analysis of POMC-derived peptides from an adult human brain sample showed comparable molar concentrations of d-α-MSH(1 −13) and β-MSH(1 −18) in both the mediobasal hypothalamus (MBH) and paraventricular hypothalamus (PVH), while β - EP(1 −31) was present at a 2- to 3-fold higher concentration. α-MSH(1-13) was 5- to 10-fold less abundant than either d-α-MSH(1-13) or β-MSH(1-18). Quantified peptide concentrations (ng/ml) were converted to nM and then normalised to the measured concentration for d-α-MSH(1-13) within each hypothalamic region.

### Human hypothalamic neurons secrete POMC-derived peptidesupon depolarisation

The secretion of POMC-derived peptides is essential for their action on downstream cells, it would
be valuable to have a human model system of this process. Since POMC-derived peptides were localised to dense core vesicle-like structures in hPSC-derived hypothalamic neurons (Fig. 2), we hypothesised that the secretion of these peptides might be enhanced by depolarisation. We therefore treated human hypothalamic cultures with either artificial cerebrospinal fluid (ACSF) or ACSF containing 30 mM KCl to induce depolarisation (Fig. 6A). By quantitative LC-MS/MS, we found that d-α-MSH(1-13), β-MSH and β-EP concentrations in culture supernatants significantly (P<0.0001) increased by 3.5- to 8.5-fold in cultures treated with KCl relative to relative to vehicle-treated cultures (Fig. 6B). These results suggest that POMC-derived peptides were trafficked into regulated secretory vesicles and released upon depolarisation.

**Figure 6:**
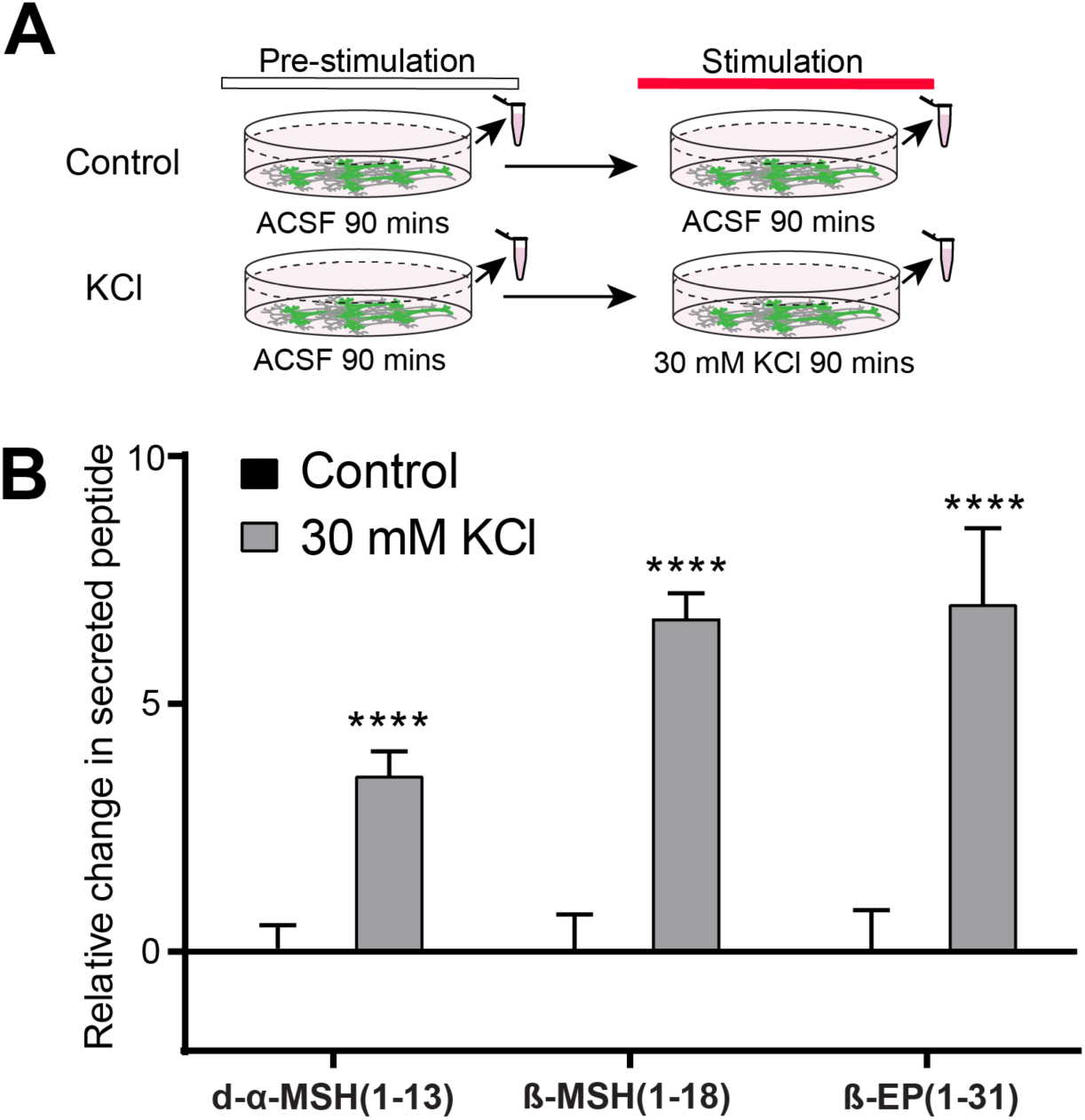
Regulated secretion of POMC-derived peptides. **A**) Experimental schematic for quantifying the stimulated secretion of POMC-derived peptides in hPSC-derived hypothalamic neurons by LC-MS/MS. Peptide concentrations measured from the supernatant during a pre-stimulation period were subtracted from the concentrations measured during control or 30 mM KCl stimulation period, converted to molarity, and normalised to control-stimulated cultures. **B**) Depolarisation of cultures with KCl significantly (P<3x10^−5^) increased the concentrations of d-α-MSH(1-13), β-MSH(1-18) and β - EP(1-31). N=3 independent experiments with 4 technical replicates per experiment. Error bars show SEM. ****, P <.0001.

### Leptin increases MSH concentration in human hypothalamic neurons

The regulation of POMC production or processing can alter the size of the pool of neuropeptides available for secretion, which could in turn modulate the downstream effects of POMC-derived peptides on energy homeostasis. Since leptin stimulates POMC neurons^54^ and induces POMC gene expression in mice^33^, we hypothesized that treatment with leptin might raise the concentration of POMC-derived peptides in human hypothalamic neurons. To test the sensitivity of the model system, we first added vehicle or a broad-spectrum inhibitor of prohormone convertases (25 µM Furin Inhibitor 1) to hPSC-derived hypothalamic neurons for 24 hours (Fig. 7A). We then added internal standards, performed quantitative LC-MS/MS, and observed a significant (P<0.01) 20-30% decrease in the measured concentration of both d-α-MSH and β-MSH in drug-treated cultures relative to vehicle controls (Fig. 7B). Next, we treated cultures with human leptin or vehicle for 24 hours and observed a significant (P<0.01) 20-30% increase in both d-α-MSH and β-MSH concentration in samples treated with 100 ng/ml (6.25 nM) leptin (Fig. 7C). These results indicate that dynamic changes in MSH concentration are detectable *in vitro* by quantitative proteomic methods, and that cultured human hypothalamic neurons respond to leptin.

**Figure 7:**
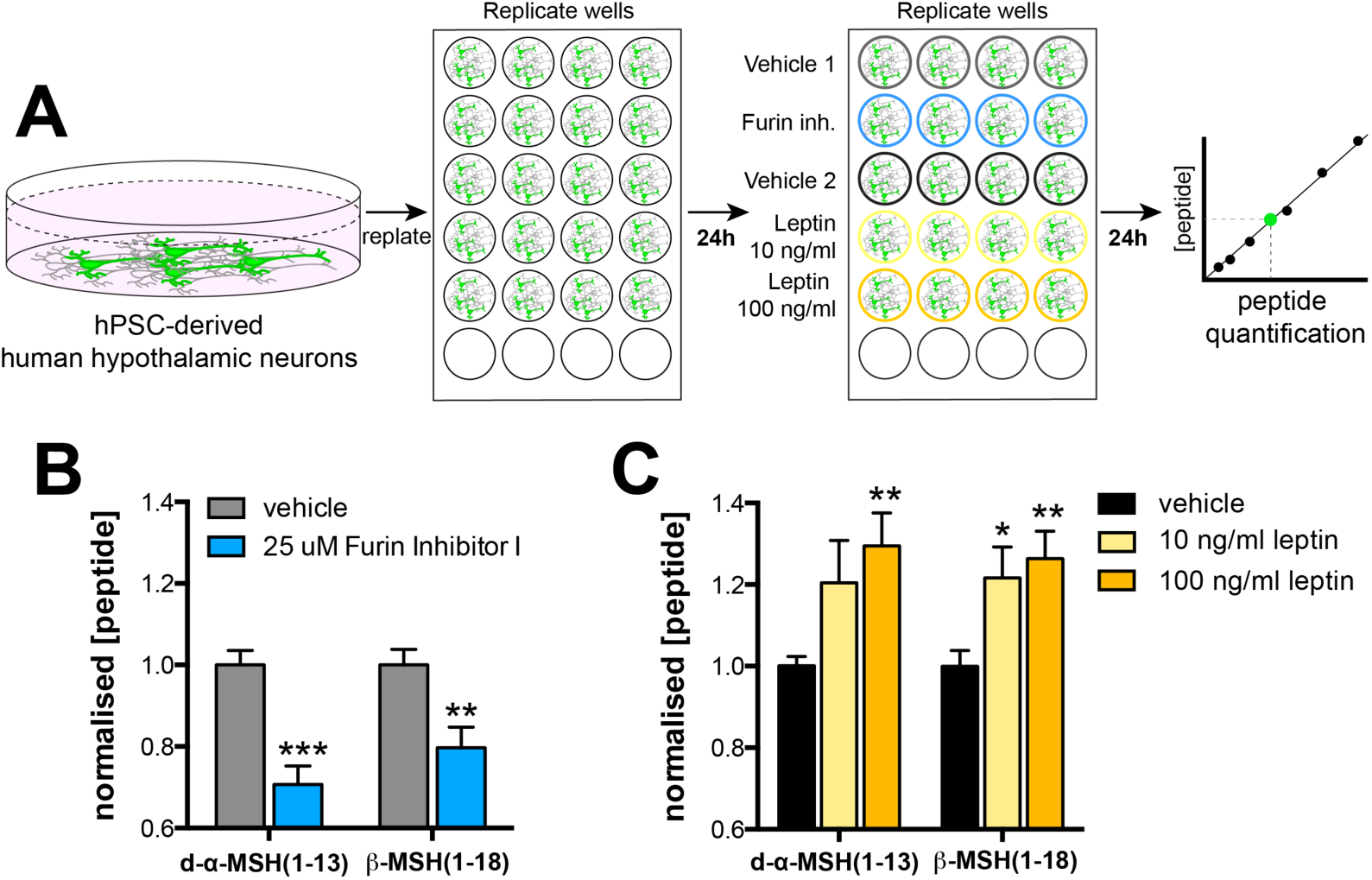
Regulation of POMC processing *in vitro*. **A**) Experimental schematic depicting the dissociation and re-plating of hPSC-derived human hypothalamic neurons for 24 hours, prior to their exposure for 24 hours to vehicle controls, 25 uM Furin Inhibitor 1 (blue), or human leptin (yellow and orange) prior to quantitative peptide analysis by LC-MS/MS. **B**) Treatment with 25 uM Furin Inhibitor 1 (blue) significantly (P <0.01) reduced measured d-α-MSH and β-MSH concentrations. **C**) Treatment with recombinant human leptin significantly (P<0.05 at 10 ng/ml, P<0.01 at 100 ng/ml) increased the measured concentrations of both d-α-MSH and β-MSH. N=2 independent experiments with 4 technical replicates per experiment. Error bars show SEM. *, P <0.05; **, P <0.01; ***, P <.001.

## DISCUSSION

In this study, we used LC-MS/MS to identify and quantify peptides in hPSC-derived neurons and in the human brain. We found that key POMC-derived peptides were correctly processed *in vitro* in all hPSC lines analysed and resembled those identified *in vivo*. On average, we observed over 2000 unique peptides per sample, including post-translational modifications. We note that these results are not exhaustive, since some larger proteins are precipitated by ACN^55^, and glycosylated peptides were not readily detected by the automated analysis pipeline we employed (see Materials and Methods). We also cannot exclude the possibility that degradation in the post-mortem interval may have affected the peptide concentrations measured in primary human brain samples. To quantify POMC-derived peptides of interest, we developed quantitative LC-MS/MS-based methods with the aid of stable isotope-labelled internal standards. These methods had comparable sensitivity and greater specificity than most ELISAs, since antibodies used in ELISAs may recognise multiple similar forms of a peptide^56^^-^^58^. Overall, our analysis revealed several noteworthy findings.

We found that β-MSH was present at substantial concentrations in both hPSC-derived neurons and in the human hypothalamus. While the 18 amino acid β-MSH sequence we observed has previously been detected in the human hypothalamus^14^, it has been suggested that this peptide and a 22 amino acid species of β-MSH might be isolation artefacts rather than true neuropeptides^19-21,59^. The use of an internal standard in our studies addressed this concern and confirmed its presence in the human brain. Several lines of evidence suggest that the importance of β-MSH in regulating energy homeostasis in the human brain has not been fully appreciated. First, it is sufficient to reduce food intake in rodents^6,60^ with similar potency to α-MSH^17^. Second, human mutations affecting either a dibasic cleavage site^15^ or a conserved tyrosine residue of β-MSH important for MC4R binding^16,17^ are strongly associated with obesity. Finally, dogs carrying frameshift mutations in POMC that affect β-MSH but not α-MSH lead to increased appetite and body weight^18^. Since β-MSH is not produced in rodents, hPSC-derived hypothalamic neurons provide a novel opportunity for studying the generation and regulation of this important peptide.

In both hPSC-derived hypothalamic neurons and in the human hypothalamus, we found that acetylated α-MSH was present at substantially lower concentrations than d-α-MSH, β-MSH, or β-EP(1-31). The use of the same internal standard for α-MSH and d-α-MSH rules out the possibility that this observation was due to poor detection, peptide degradation, or a systematic technical error in quantification. In agreement with our results, d-α-MSH was previously shown to be more abundant than α-MSH in the rodent hypothalamus^35,61-64^ and human brain^65^. Since peptide degradation in the post-mortem interval is more likely to affect d-α-MSH than the relatively stable α-MSH^35^, our *in vivo* results may underestimate the relative prevalence of d-α-MSH in the human brain. The biological significance of this finding is unclear, since we cannot exclude the possibility that only a small proportion of the d-α-MSH pool is acetylated before secretion^8^, and that acetylated α-MSH (and perhaps also β-MSH) may be the primary ligand of MC4R *in vivo*. However, it is likely that at least some d-α-MSH is secreted, raising the question of what its role might be in human energy homeostasis. Previous studies using acute peptide injections suggested that d-α-MSH is less potent at reducing food intake in rodents than α-MSH^6,52^. These findings have recently been challenged by a longer-term study in which both α-MSH and d-α-MSH were genetically deleted from mice, and then individually infused for 14 days at physiological levels into the mouse brain. This study revealed that both peptides could similarly reduce appetite and body weight in male mice^5^. Furthermore, d-α-MSH has been reported to be a more potent agonist of MC4R than α-MSH^8^. We therefore suggest that in humans, the relative importance of both d-α-MSH and β-MSH may have been underestimated, and that the assumption that acetylated α-MSH is the principal anorexigenic melanocortin peptide in humans should be reconsidered.

Whereas we found that d-α-MSH and β-MSH were present at roughly equimolar concentrations both *in vitro* and *in vivo*, we observed b-EP(1-31) at 3-4 fold higher concentrations in hPSC-derived neurons and 2-3 fold higher concentrations in the brain. The use of stable-isotope labelled internal standards makes it unlikely that this discrepancy could be explained by peptide oxidation during the extraction process. However, we cannot exclude the possibility that peptide standards might have differential stability when stored at −80°C. Another potential explanation is that β-EP(1-31) might be more readily produced in hypothalamic neurons as it only requires a single cleavage event at a Lys-Arg 219-220 (position 235-236 from translational start) to be liberated from larger precursor peptides, whereas the generation of either d-α-MSH or β-MSH requires at least two cleavage events.

Predictions made by studying hypothalamic neurons derived from three distinct hPSC lines were largely confirmed in primary human hypothalamic samples. However, found that although α-MSH was consistently less abundant than either d-α-MSH or β-MSH, this difference was less pronounced *in vivo*. Furthermore, the human brain contained a larger repertoire of neuropeptides than we observed *in vitro*. Some of these differences might be explained by the distinct anatomical origin of neuropeptide-producing neurons, since our *in vitro* model generates cells with a predominantly ventral hypothalamic regional identity^66^. For example, neurons producing gonadotropin releasing hormone (GNRH) are generated in the olfactory placode and then migrate into the hypothalamus^67^, and the glucagon like peptide 1 (GLP-1) we detected *in vivo* was likely present in the axon terminals of proglucagon-expressing neurons whose cell bodies are found in the nucleus of the solitary tract (NTS) in the brainstem in rodents^68^. We do not expect either of these cell types to be produced when hPSCs are patterned to ventral hypothalamic cells. Other differences could be explained by the relative immaturity of hPSC-derived hypothalamic neurons at the time points we studied, resulting in incomplete pro-peptide processing or expression at levels too low to be automatically detectable using the automated peptide identification pipeline we employed. The use of more sensitive LC-MS/MS instruments or the analysis of raw peptidomic data using new discovery tools^69^ may reveal further biologically important peptides.

We found that exposure of hPSC-derived hypothalamic neurons to leptin significantly increased the concentration of both α-MSH and β-MSH, and that these peptides were robustly secreted from upon stimulation. These findings are consistent with observations in animal models, suggesting that other discoveries made using this human cellular model system may have predictive power for understanding human-specific aspects of POMC biology, such as the regulation of β-MSH processing and secretion. The genetic or environmental manipulation of hPSC-derived hypothalamic neurons coupled with quantitative LC-MS/MS-based analysis may provide insights into the molecular mechanisms regulating POMC processing and identify novel therapeutic targets for obesity.

## MATERIALS AND METHODS

### Differentiation of hPSCs to hypothalamic and cortical neurons

Pluripotent stem cell maintenance, hypothalamic differentiation^36,37^, and cortical differentiation^70^ was carried out as previously described. Cell lines used were WA09(H9) hESCs (Passage 38-42, WiCell, RRID: CVCL_9773), HUES9 hESCs (Passage 32-40, Harvard University, RRID: CVCL_0057), and FSPS13B hiPSCs (Passage 47-52, a kind gift from L. Vallier). Briefly, hPSCs were maintained and expanded in mTeSR1 media (Stem Cell Technologies, Cat# 85850). For hypothalamic differentiation, hPSCs were grown in N2B27 media and treated from day 0-10 with 10 µM SB431542 (Sigma-Aldrich, Cat# S4317), 100 nM LDN193189 (Stemgent, Cat# 04-0074) 2 µM XAV939 (Stemgent, Cat# 04-0046), in combination with agonists of Sonic Hedgehog pathway 1 µM Purmorphamine (Calbiochem, Cat# 540220) and 1 ?M SAG (Fisher Scientific, Cat# 56-666) on days 2-8, and then followed 5 µM DAPT (Sigma-Aldrich, Cat# D5942) from day 8-14. Cells were then maintained in N2B27 medium supplemented with 10 ng/mL BDNF (Peprotech, Cat# 450-02). Experiments were carried out on hypothalamic cultures 25-90 days post-differentiation, at which point POMC could be robustly detected by immunohistochemistry. Cultures were maintained in 24-well plates (Corning, Cat# 3524) for proteomic experiments, in 8-well Ibidi dishes (Thistle Scientific, Cat# IB-80826) for confocal imaging, and in Labtek 4-well chamber slides (Sigma-Aldrich, Cat# C6932) for Immunogold EM.

### Immunocytochemistry and confocal imaging

Cells were fixed in 4% w/v paraformaldehyde in PBS for 10 minutes at room temperature. After three washes in TBS, cultures were incubated overnight at 4°C with primary antibody diluted in 10% normal donkey serum in TBS with 0.1% Triton X-100. Primary antibodies used were raised against MAP2 (1:2,000, Abcam, Cat# ab5392, RRID:AB_2138153), TUJ1 (TUBB3) (1:2,000, BioLegend, Cat# 845502, RRID:AB_2566589), or POMC (A1H5, A3H9, N1C11, used at 1:5,000, developed by Prof. Anne White). Primary antibody was removed by washing three times in TBS. Cells were then incubated in Alexa Fluor-conjugated secondary antibodies (Thermo Fisher Scientific) diluted 1:500 in 10% normal donkey serum in TBS with 0.1% Triton X-100 for 2 hours at room temperature. After three washes in TBS, cells were incubated in 360 nM DAPI in TBS, and then washed a further three times in TBS. Cells were imaged in TBS on a Zeiss LSM 510 confocal microscope. Images were analysed and processed with ImageJ (NIH, RRID:SCR_003070).

### Immunogold labelling

hiPSC-derived hypothalamic neurons were grown in 4-well Nunc Lab-Tek chamber slides (Sigma-Aldrich, Cat# C6932) as described above. Cells were fixed in 4% paraformaldehyde and 0.5% glutaraldehyde in 0.1 M phosphate buffer (PB) for 1 hour at 37°C. Pre-embedding Immunogold labelling was performed by incubation with either mouse anti POMC A1H5 (1:2,500; developed by Prof. Anne White) or rabbit anti β-endorphin (1:500; Millipore, Cat# Ab5028, RRID:AB_91645) primary antibodies as previously described^71^.

### Electron microscopy

Cells were postfixed in 1% osmium tetroxide, 7% sucrose in 0.1 M PB for 30 minutes at room temperature, washed in deionized water, and partially dehydrated in ethanol. Cells were then stained in 2% uranyl acetate in 70% ethanol in the dark for 150 minutes at 4°C. Cells were further dehydrated in ethanol, and infiltrated overnight in Durcupan ACM epoxy resin (Fluka, Sigma-Aldrich, St. Louis, USA). Following resin hardening, embedded cell cultures were detached from the chamber slide and glued to resin blocks. Serial semi-thin sections (1.5 µm) were cut with an Ultracut UC-6 ultramicrotome (Leica, Heidelberg, Germany) and mounted onto glass microscope slides and lightly stained with 1% toluidine blue. Selected semi-thin sections were glued with Super Glue-3, Loctite (Henkel, Düsseldorf, Germany) to resin blocks and subsequently detached from the glass-slides by repeated freezing (in liquid nitrogen) and thawing. Ultra-thin sections (70-80 nm) were obtained with the ultramicrotome from detached semi-thin sections, and further stained with lead citrate (Reynolds’ solution)^72^. Finally, cells were imaged at 80 kV on a FEI Tecnai G^2^ Spirit transmission electron microscope (FEI Europe, Eindhoven, Netherlands) equipped with a Morada CCD digital camera (Olympus Soft Image Solutions GmbH, Mϋnster, Germany).

### Tissue homogenization and peptide extraction from cell pellets and ACSF

To extract and solubilise peptides, cultures were partially dissociated by treatment with 1 mM EDTA in PBS for 10 minutes. EDTA was then removed and cells were directly lysed by vigorously pipetting cells with 80% acetonitrile (ACN, Pierce Cat# 51101), an organic solvent that is routinely used to precipitate larger proteins and protein complexes^53,73^. This lysate was then transferred to low protein-binding microcentrifuge tube (Sigma, Cat# Z666505), vortexed for 1 minute, and pelleted for 15 minutes at 12,000 *g* at 4°C. The supernatant was then transferred to a new low protein-binding microcentrifuge tube or 96-well low protein-binding plate (VWR, Cat# 951032921) and immediately processed for analysis (see below) or stored at −80°C.

In some experiments, cells were lysed in 6M guanidine hydrochloride (GuHCl)^74,75^ (Sigma, Cat# G4505) and transferred to low-protein-binding microcentrifuge tubes (Sigma, Cat# Z666505) (Personal communication, Pierre Larraufie et al.). Lysates were snap-frozen on dry ice and thawed on ice three times to aid with lysis, before four parts 80% acetonitrile were added and mixed thoroughly. Phases were separated by centrifugation for 5 minutes at 12,000 *g* at 4°C. The bottom aqueous phase was transferred to a fresh low protein-binding microcentrifuge tube, or a 96-well low protein-binding plate (VWR, Cat# 951032921) and immediately processed for analysis (see below) or stored at −80°C.

Samples processed with 80% ACN or GuHCl-ACN were thawed on ice if previously frozen. Samples were either dehydrated in an evaporating centrifuge or evaporated with pure nitrogen at 40°C on an SPE Dry evaporator system (Biotage, Uppsalla, Sweden). Dehydrated peptides were reconstituted with 0.1% formic acid in dH_2_O, spiked with 500 pg stable-isotope internal standards for d-α-MSH(1-13), β-MSH(1-18) and β-EP(1-31), and loaded onto a HLB Prime micro elution plate (Waters, Cat# 186008052) for solid-phase extraction. Plates were washed with 200 µL 0.1% formic acid in dH_2_O (v/v), followed by a wash with 200 µL 1% acetic acid with 5% methanol in dH_2_O (v/v/v). Samples were eluted with 2x 30 µL of 60% Methanol in dH_2_O with 10% Acetic acid (v/v) and diluted further with 75 µL of 0.1% formic acid in dH_2_O (v/v) and stored at −80 °C until loading onto LC columns for MS/MS.

ACSF from stimulation experiments was collected and immediately snap-frozen on dry ice and stored at −80°C. Samples were then thawed on ice, followed by spike-in of 0.5 ng stable-isotope internal standards for α-MSH(1-13), d-α-MSH(1-13), β-MSH(1-18) and direct solid-phase extraction, as described above.

### Stimulated secretion of POMC-derived peptides

hPSC-derived neurons were washed three times with HEPES-buffered ACSF consisting of 129 mM NaCl, 5 mM KCl, 1 mM CaCl_2_, 1 mM MgCl_2_, 25 mM HEPES, 11.1 mM Glucose. Cells were incubated in ACSF for 90 minutes at 37°C. The supernatant was collected from the cells and snap-frozen on dry ice. After removing the ACSF, the cells were incubated for 90 minutes at 37°C in either ACSF as described above or a 30 mM KCl solution consisting of 104 mM NaCl, 30 mM KCl, 1 mM CaCl_2_, 1mM MgCl_2_, 25 mM HEPES, 11.1 mM Glucose, or a 129mM KCl solution consisting of 5 mM NaCl, 129 mM KCl, 1 mM CaCl_2_, 1mM MgCl_2_, 25 mM HEPES, 11.1 mM Glucose. Media supernatants from control ACSF or 30 mM KCl were collected, snap-frozen, and stored at −80°C. To account for slight well-to-well variation in the amount of POMC neurons or POMC-derived peptide production, KCl-induced secretion during the stimulation period (Fig. 5A) was normalized by subtracting the quantified pre-stimulation value for each POMC-derived peptide from the post-stimulation value.

### Regulation of MSH concentration by exogenous factors

hPSC-derived hypothalamic neuronal cultures were dissociated after D25, at which point POMC neurons could clearly be visualised by immunohistochemistry, and re-plated in parallel cultures as previously described^37^. After 24 hours of recovery, media was changed and culture were treated for 24 hours with experimental agents. These included leptin (PeproTech, Cat# 300-27) reconstituted in PBS+0.1% BSA to a final concentration of 10 ng/ml or 100 ng/ml or vehicle (PBS+0.1% BSA), and Furin Inhibitor 1 (Cayman Chemical Co., Cat# B00165) added to a final concentration of 25 µM or vehicle (DMSO to 0.1% final concentration). Cultures were then processed for LC-MS/MS as described above.

### Peptide discovery by LC-MS/MS

Peptide extracts were run using nano-flow-based separation and electrospray approaches on a Thermo Fisher Ultimate 3000 nano-LC system coupled to a Q Exactive Plus Orbitrap mass spectrometer (ThermoScientific, San Jose, USA) as described previously^53^. Extracts were injected at a flow rate of 30 µL/minute onto a peptide trap column (0.3 × 5 mm; ThermoFisher Scientific), washed for 15 minutes, and switched in line with a nano-easy column (0.075 × 250 mm; ThermoFisher Scientific) flowing at 300 nL/minute. Both nano and trap column temperatures were set at 45°C during the analysis. The buffers used for nano-LC separations were A: 0.1% formic acid in water (v/v) and B: 0.1% formic acid (v/v) in 80:20 ACN/water. Initial starting conditions were 2.5% B (equating to 2% ACN), and held for 5 minutes. A ramp to 50% B was performed over 135 minutes, followed by a wash with 90% B for 20 minutes before returning to starting conditions for 20 minutes, totalling an entire run time of 190 minutes. Electrospray analysis was performed using a spray voltage of 1.8 kV, the tune settings for the MS used an S-lens setting of 70 v to target peptides of higher m/z values. A full scan range of 400-1600 m/z was used at a resolution of 75,000 before the top 10 ions of each spectrum were selected for MS/MS analysis. Existing ions selected for fragmentation were added to an exclusion list for 30 s.

### Peptide identification using Peaks

The acquired LC-MS files were searched using Peaks 8.5 software (Waterloo, ON, Canada) against the human Swissprot database (06-May-2016, 26-October-2017). A no-digest setting was used, which enabled peptides of up to 65 amino acids in length to be matched, and precursor and product ion tolerances were set at 10 ppm and 0.05 Da, respectively. A fixed post-translational modification of carbamidomethylation was applied to cysteine residues, whilst variable modifications included methionine oxidation, N-terminal pyro-glutamate, N-terminal acetylation and C-terminal amidation. A minimum of 1 unique peptide and a false discovery rate value of 1% was used to filter the results.

### Peptide quantification by LC-MS/MS

Calibration standards of synthetically produced α-MSH(1-13), d-α-MSH(1-13), β-MSH(1-18) and β-EP(1-31) (Bachem, Bebendorf, Switzerland) to 0.01-10 ng/mL in a matrix of 0.001% BSA in 0.1% formic acid (v/v), or in lysates from cultured cortical neurons that contain no detectable POMC peptides. Calibration standards were then spiked with 0.5 ng custom-ordered stable-isotope internal standards (Cambridge Research Biochemical Ltd.) for:

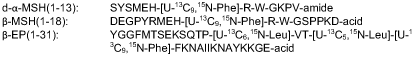

Samples were extracted using the solid-phase-extraction method described above. LC-MS instrumentation used for the quantitation of the POMC-derived peptides included an H-Class Acquity (Waters) attached to a TQ-XS triple quadrupole mass spectrometer (Waters). Sample (10 µL) was injected onto a 2.1 x 50 mm 1.8 µm particle HSS T3 Aquity column held at 60 °C and flowing at 700 µL/minute. Gradient starting condition were 87.5 % A (0.1% formic acid in water v/v) and 12.5 % B (0.1% formic acid in ACN). Starting conditions were held for 30 seconds before raising to 30 % B over 2.5 minutes. The column was flushed with 90 % B for 1.1 minutes before returning to starting conditions. The total time of each analysis was 5 minutes, with the first 1.1 minute and last 1.8 minute diverted to waste. Mass spectrometry conditions involved targeting four POMC-derived peptides, and the transition and collision energy details for each peptide (along with associated internal standards are given below:

| Peptide | Q1 (m/z) | Q3 (m/z) | Collision energy (eV) | Dwell time (ms) |
| --- | --- | --- | --- | --- |
| Des-Ac-α-MSH | 406.55 | 458.13 | 12 | 40 |
| Des-Ac-α-MSH IS | 409.05 | 461.56 | 12 | 40 |
| Ac-α-MSH | 555.98 | 686.75 | 19 | 40 |
| β-MSH | 552.1 | 654.08 | 20 | 40 |
| β-MSH IS | 554.6 | 567.41 | 20 | 40 |
| β-Endorphin | 693.62 | 825.89 | 22 | 40 |
| β-Endorphin IS | 698.52 | 832.19 | 22 | 40 |

The source parameters used included a positive electrospray ion spray voltage of 3.0 kV, gas flow of 600 L/hour, desolvation temperature of 500 °C and a cone voltage of 40 V. Peptide peak areas were integrated using the TargetLynx program associated with Masslynx V 4.2 (Waters) and peptide peak areas were ratiometrically compared against their corresponding stable isotope-labelled internal standard peptide. The POMC-derived peptide peak area ratio concentrations in extracted samples were compared against the calibration lines to assign concentration values. The peptide calibration lines showed consistent results over a thousandfold range of peptide concentrations (R^2^>0.98 for all peptides).

### Human brain samples

Fresh human brain samples were obtained from two donors under informed consent via the Cambridge Brain Bank after ethical committee approval. The first donor was a 75-year-old woman who died from kidney cancer. The second donor was an 81-year-old woman who died from leukaemia and suffered from chronic kidney disease. The total post-mortem intervals for these two donors were 24 hours and 57 hours, respectively. Neither donor had a history of neurodegenerative disease, nor did the brains have obvious pathological evidence of neurodegeneration. After death, brains were dissected from the skull and stored in buffered saline at 4?C until the time of dissection for LC-MS/MS. Dissected brain regions of approximately 3x3x3 mm encompassed the mediobasal hypothalamus (MBH) that should contain POMC cell bodies, the paraventricular nucleus of the hypothalamus (PVH) that should contain POMC neuron axon terminals, and a small piece of the orbitofrontal gyrus of the cerebral cortex (CTX) extending through all cortical layers. Immediately after dissection, primary samples were disrupted in 6M GuHCl at 4°C in Lysing Matrix D tubes (MPbio, 116913100) to stabilise and extract peptides, and then fully homogenized using a Fastprep 24 5G homogeniser (MPbio) using four 40 second runs at 6 m/s separated by 5 minutes of cooling on ice between each homogenisation cycle. Samples were stored at −80°C or processed immediately. Peptides were extracted as described above and analysed by nanoflow LC-MS/MS.

### Statistical analysis

Statistical analyses in Figure 4D, 4E, 6B, 7B, 7C, S1C, S1E were carried out using Prism (GraphPad, RRID:SCR_002798). Significance was calculated using the Holm-Sidak method, with alpha 0.05. Error bars in Figure 4D, 4E, 6B, 7B, 7C, S1C, S1E show the standard error of the mean (SEM).

## ACKNOWLEDGEMENTS

We thank the brain donors and their families for their generosity, as well as Jenny Wilson, Dr. Poonam Singh, Dr. Kathryn Brown, and Dr. Robert Fincham and others at the Cambridge Brain Bank (CBB) for assisting with human sample procurement and delivery, and Dr Kieren Allinson for assisting with the dissection of human hypothalamic nuclei. The CBB is supported by the NIHR Cambridge Biomedical Research Centre. We also thank Phil Lowry and Tony Coll for their comments that improved this manuscript. This work was supported by the Academy of Medical Sciences (SBF001\1016), Medical Research Council (MR/P501967/1, MR/M009041/1, and Metabolic Diseases Unit, award 4050281695, MC UU 12012/3, MC UU 12012/5,), Wellcome Trust (106262/Z/14/Z, 106263/Z/14/Z, PSAG/097), Prometeo network (PROMETEOII/2014/075), Red de Terapia Celular TerCel of the Instituto de Salud Carlos III (RD16/0011/0026), Mawer-Fitzgerald endowment, Manchester Academic Health Sciences Centre, and by a project grant from MedImmune.

## AUTHOR CONTRIBUTIONS

PK and FTM conceived of the project, analysed data, and wrote the manuscript with input from all co-authors. PK performed hypothalamic differentiation with the help of MJ and BB, performed experiments, and prepared samples for LC-MS/MS. RK extracted and ran samples for LC-MS/MS and assisted with data analysis. PL helped develop methods for peptide extraction and assisted with sample preparation. VHP generated and helped interpret electron microscopic data. AW contributed anti-POMC antibodies and the POMC ELISA protocol. TB performed the POMC ELISA. JP helped develop the quantitative LC-MS/MS SRM methodology and analysed human samples. FG, FR, ISF, and SOR financially supported this study and provided intellectual guidance.

